# A new family of small ArdA proteins reveals an antirestriction activity

**DOI:** 10.1101/2025.03.20.644391

**Authors:** A.A. Utkina, A.A. Kudryavtseva, O.E. Melkina, S.M. Rastorguev, A.V. Vlasov, K.S. Pustovoit, I.V. Manukhov

## Abstract

Antirestriction proteins are known to protect mobile genetic elements from the host’s restriction-modification (RM) systems. In our study, we identified a new family of small proteins, which we named sArdA The sArdA proteins are homologous to DNA-mimicking ArdA proteins but differ in size, being approximately one-third the length of full ArdAs. Moreover, sArdA family contains two subgroups, one of which is structurally similar to the N-terminal end of ArdA, while the other one – to the C-terminal end. Phylogenetic analysis demonstrated that genes encoding these proteins evolved into evolutionarily stable subfamilies, named sArdN and sArdC, respectively. Alphafold structure prediction of sArdA interaction with RM systems revealed four states of EcoKI, which differ the angle between its two M-subunits while interacting with different agents. Interestingly, both sArdN and sArdC triggered the same intermediate closed state of EcoKI indicating the possible new interaction pathways of Ards with RM systems.

For phenotypic studies in *Escherichia coli* cells, we cloned the *sardN* gene from the chromosome of *Corynebacterium pilbarense* and the *sardC* gene from *Lactococcus cremoris*. Both genes were shown to protect λ phage DNA from restriction by the type I RM system. However, they revealed specificity to different restriction-modification systems. Specifically, *sArdC* was more effective against EcoR124II, whereas *sArdN* was more potent against EcoKI. Furthermore, both genes demonstrated high antimethylation activity against EcoKI. Our current findings suggest the idea that binding specificity of DNA-mimicking proteins to their targets could also be achieved by very short proteins.

## Introduction

Type I RM systems, consisting of restriction endonucleases and DNA methyltransferases, serve as a critical defense mechanism in bacteria by recognizing and cleaving foreign DNA (Murray, 2000). ArdA proteins can counteract this defense by mimicking the structure and surface charge of DNA, thereby preventing the degradation of the mobile genetic element’s DNA (Tock and Dryden, 2005). *ard*A genes are commonly found within conjugative plasmids, transposons, and bacteriophage DNA and are typically among the first genes to enter the cell during horizontal gene transfer (Chilley and Wilkins, 1995).

Recent evidence shows chromosomal ArdA protein from *Bifidobacterium bifidum* modulating bacterial gene expression in *E. coli* (M. V. Gladysheva-Azgari et al., 2023) raising intriguing questions about their cellular roles.

Here, we identify two novel ArdA-like antirestriction proteins from *L. cremoris* (further named ArdA_1576) and *C. pilbarense* (further named ArdA_8247). These newly identified proteins are particularly intriguing due to their exceptionally small sizes, being encoded by genes of 342 and 264 base pairs respectively, making them some of the smallest ArdA proteins discovered to date.

## Materials and Methods

### Bacterial Strains and Plasmids

*E. coli* TG1 (K-12 glnV44 thi-1 Δ(lac-proAB) Δ(mcrB-hsdSM)5(rK–mK–) F′ [traD36 proAB+ lacIq lacZΔM15]) strain was used as the host strain for all experiments. Two novel, exceptionally small chromosomal *ardA* genes were cloned from the *C. pilbarense* B-8247 and *L. cremoris* B-1576 chromosomes. All strains were obtained from the All-Russian Collection of Industrial Microorganisms (VKPM).

### Construction of Plasmids

The two novel chromosomal *ardA* genes were amplified from *C. pilbarense* and *L. cremoris* genomes via PCR using primers incorporating NdeI and EcoRI restriction sites. The PCR products were purified and digested with NdeI and EcoRI restriction enzymes. The pIR-DPAl expression vector was also digested with the same restriction enzymes. The digested *ardA* gene fragments and the linearized pIR-DPAl vector were ligated using T4 DNA ligase (Thermo Fisher Scientific, USA). The ligation products were transformed into calcium-competent *E. coli* cells, and transformants were selected on LB agar plates containing kanamycin (50 μg/mL). Plasmids were isolated from positive colonies and confirmed by sequencing. The resulting plasmids (Table 1) contained the target *ardA* genes under the control of the pIR-DPAl expression vector promoter.

### Antirestriction activity assays

To evaluate the antirestriction activity of ArdA_8247 and ArdA_1576 against the EcoKI and EcoR124II restriction-modification (RM) systems, unmodified lambda phage (λ_0_) was used. Three types of *E. coli* TG1 cultures were used: cells without plasmids, cells harboring plasmids containing the genes for the RM systems, and cells carrying plasmids with the RM system genes along with either *ardA_8247* or *ardA_1576* (as detailed in Table 1). The plasmid pIR-DPAl-ArdA-R64, containing a well-described *ardA* gene from the conjugative plasmid R64, served as a positive control. Phage plating and calculation of the efficiency of plating (EOP) were conducted following established procedures (Kudryavtseva et al., 2023b).

The efficiency of plating (EOP) was estimated as

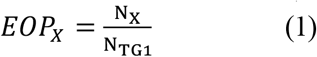

where N_X_- the number of λ.0 phage plaques on the *E. coli* TG1 cells carrying genes ‘X’ affecting the plaque forming, N_TG1_ – the number of λ_0_ phage plaques on *E. coli* TG1 (without any additional restriction or antirestriction genes).

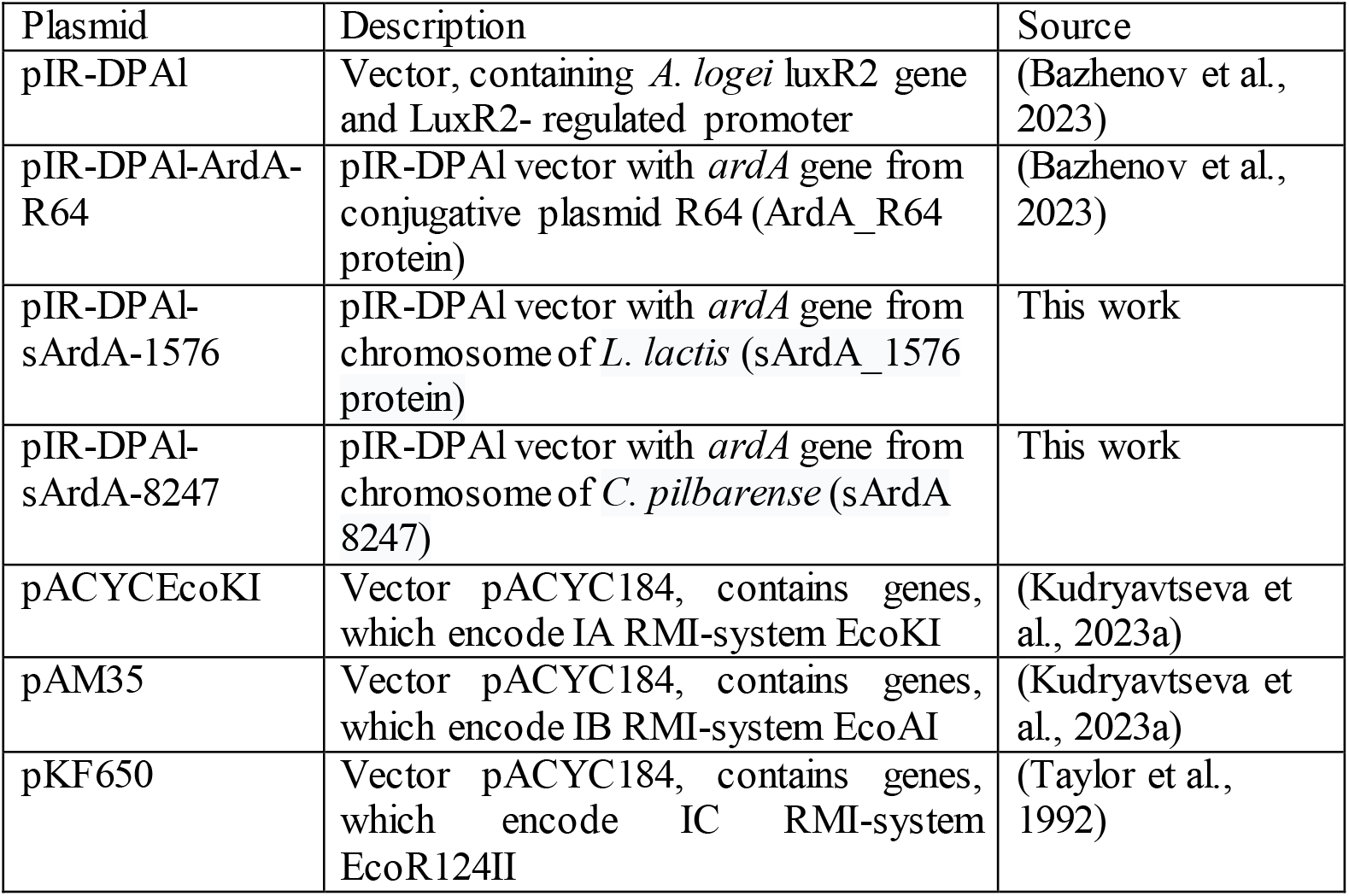

### Phylogenetic studies

The protein sequences from the NCBI database used to construct phylogenetic trees are presented in Table S1. We used ModelFinder (Kalyaanamoorthy et al., 2017) to select the best evolutionary model for our sequence alignments. The phylogenetic relationships between ArdA proteins were visualized using the Tree Of Life (iTOL) v5 online tool (Letunic and Bork, 2021).

### Alphafold structure predictions

We performed structure predictions of the proteins sArdA_1576, sArdA_8247, ArdA_916 and their interaction with the EcoKI (S_1_M_2_) protein complex using the AlphaFold v.3 software (Abramson et al., 2024). For reference, prediction of the interaction between the EcoKI (S_1_M_2_) protein complex and dsDNA (AAAACACGTGTGTGCAA) was performed.

### Antimethylation activity assay

To test the antimethylation activity of newly described sArdA proteins we used *E. coli* AB1157 [*thr-J leu-6 proA2 his4 thi-J argE3 lacY galK2 ara-14 xyl-5 mtl-l tsx-33 rpsL31* supE44] strain which contains EcoKI RMI system. We collected λ_k_ from AB1157 strain and λ_k1576_ and λ_k8247_ from AB1157 pIR-DPAl-ArdA-1576 and AB1157 pIR-DPAl-ArdA-8247 respectively. After that the EOP was estimated.

## Results

Recently we showed that *ardA* genes can be found inside bacterial chromosomes (M. V Gladysheva-Azgari et al., 2023),(Kudryavtseva et al., 2024). We searched for *ardA*-like genes across various bacterial species and identified a new group of chromosomal *ardA* genes which are twice smaller than classic *ardA* genes such as *ardA* from Tn_916 transposon (McMahon et al., 2009). It should be noted that these small genes (CDS) are not surrounded be the *ardA* gene fragments, within bacterial chromosomes, but seems to be evolving by themselves. We named this group of ArdA proteins “small ArdA” (sArdA) and performed a protein alignment using T-coffee service (Notredame et al., 2000). The alignment results were visualized via MView web service at EMBL-EBI site (Madeira et al., 2019) (figure 1).

**Figure 1.**
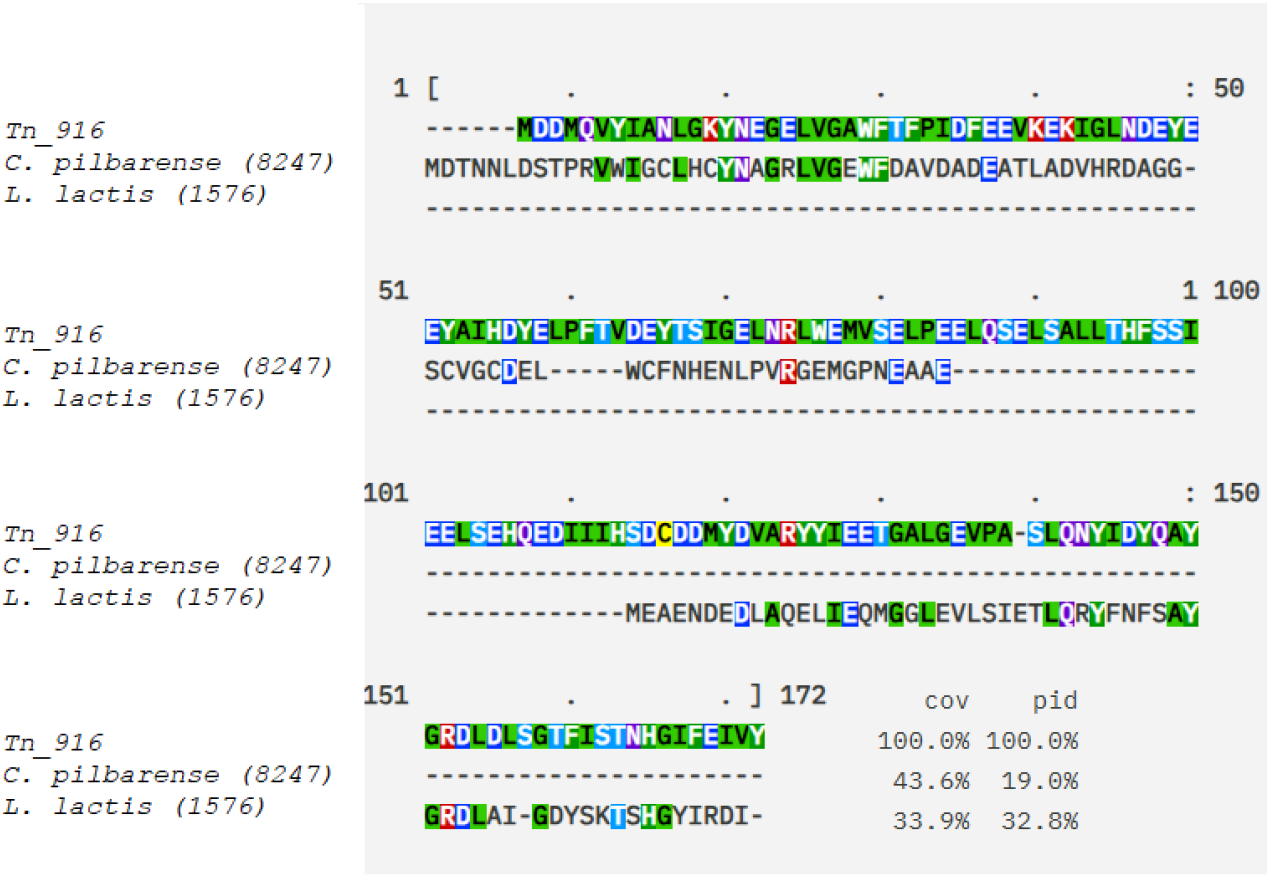
Alignment of the amino acid sequences encoded by the *sardA* genes from *C. pilbarense* and *L. lactis* chromosomes. All proteins are aligned to the canonical ArdA protein from Tn916 transposon. Designations: (–) – no homologous residue; cov – coverage; pid – percent identity.

Figure 1 illustrates that although the percentage of amino acid identity is rather low, a distinctive negative charge (indicated by negatively charged amino acids colored in blue), mimicking DNA structure, is preserved throughout all aligned protein sequences. Subsequently, we generated structural models of the selected proteins, sArdA_1576 and sArdA_8247, using the AlphaFold3 service(Abramson et al., 2024). These modeled structures were subsequently aligned with the canonical rdA_Tn916 structure(McMahon et al., 2009).

From the Figure 2, it is evident that despite the low percentage of identical amino acids (19%), there exists a structural similarity between sArdA_8247 and the N-terminal region of the classical full-length ArdA from Tn916, as well as with the the C-terminal segment of ArdA aligns to the studied protein sArdA_1576. Moreover, sArdA_1576 aligns not just to the C-terminal region but specifically with the dimerization interface of the full-length ArdA_Tn916, thereby overlapping the dimerization site of its subunits.

**Figure 2.**
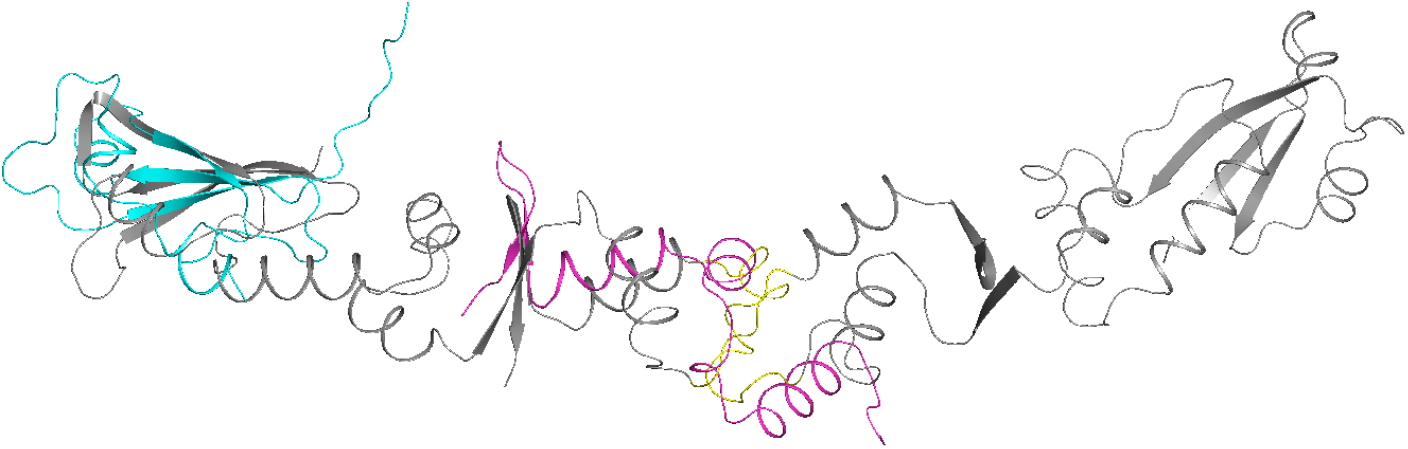
Structural alignment. Gray — dimeric structure of ArdA_Tn916, yellow highlights amino acids forming the interaction interface between two ArdA_Tn916 subunits (‘antirestriction motif’(Rastorguev and Zavilgelsky, 2003)). Blue — predicted structure of sArdA_8247, pink — predicted structure of sArdA_1576.

Then, we searched for a short *sardA* genes which are homologous to the N- and C-terminal regions of the full-length *ardA* gene across various microorganisms. The results of phylogenetic analysis with the identified representatives of *sardA* are presented in Figure 3. Homologues to the N-terminus are henceforth referred to as sArdN, and those to the C-terminus as sArdC.

**Figure 3.**
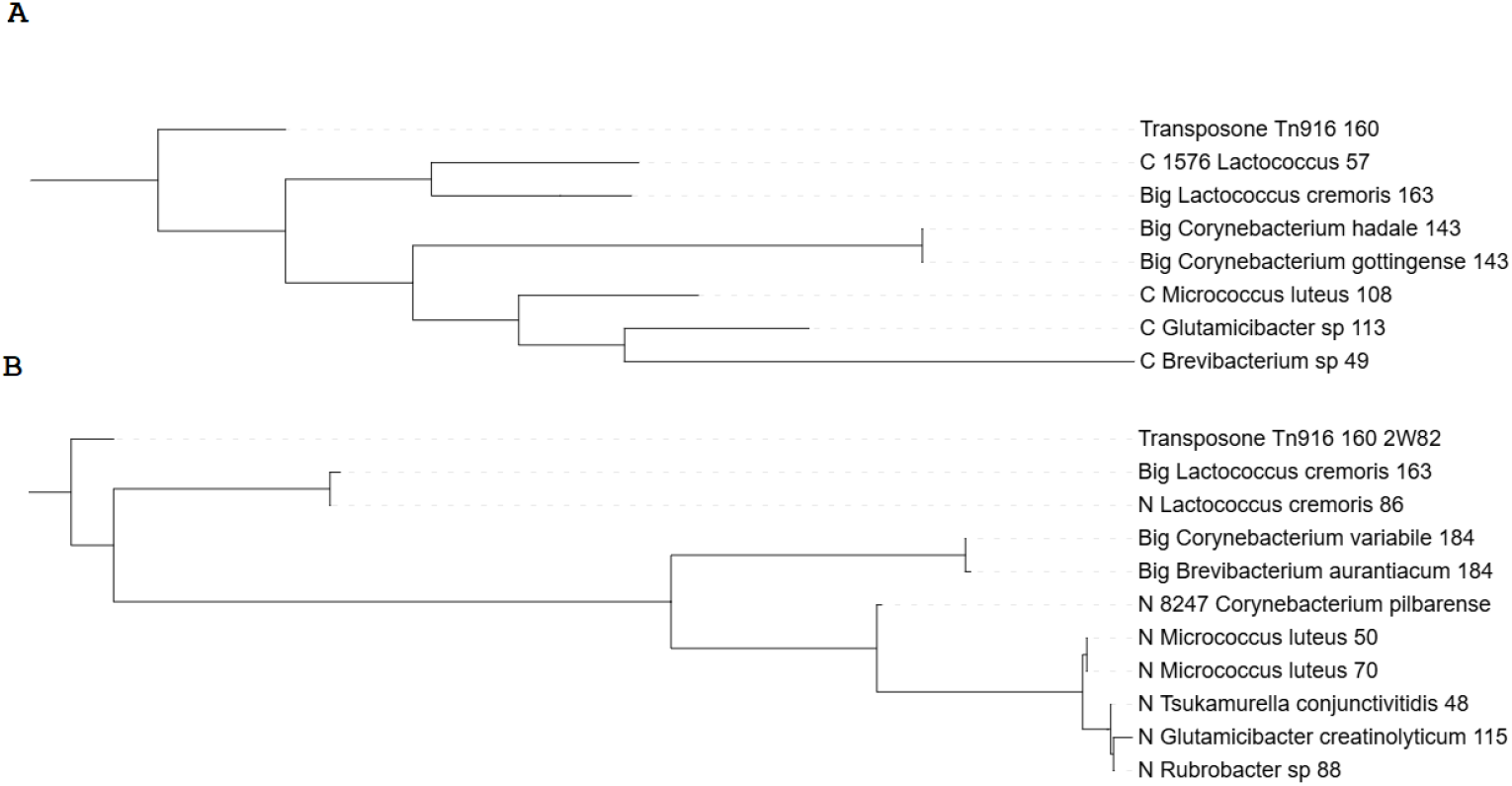
Phylogenetic trees for sArdA proteins from bacterial chromosomes from genus *Lactococcus* and phylum *Actinomicetes*. ArdA_Tn916 sequence was used as an outgroup. The ‘N’ before the species name means that the protein aligns to the N-terminus, the ‘C’ means that the protein aligns to the C-terminus, Big – means ‘full’-sized ArdB. The numbers at the end indicate the protein length in aa.

From the data shown in Figure 3, it is clear that small representatives of the *ardA* gene family are widely distributed among various bacterial species. Specifically, within the phylum *Actinomicetes* sArdA proteins cluster into two distinct subgroups—subfamilies. We have provisionally named these subfamilies sArdN (homologs of the N-terminal region) and sArdC (homologs of the C-terminal region of the classical ArdA protein), as depicted in Figures 3A and 3B respectively. The clustering of sArdA across diverse bacterial species shows that, once established through evolutionary processes, these genes continue to be preserved, forming clusters of conserved sequences.

We note that both sArdN and sArdC from the genus *Lactococcus* cluster with the full-length ArdA protein also presented in these bacteria, rather than with their respective subfamilies from other taxa belonging to *Actinomicetes*. This suggests that the small *ardA* genes likely arose multiple times during evolution.

To validate our results, we performed structure predictions of EcoKI(S _1_M_2_, one S-subunit, two M subunits) interacting with sArdC and sArdN using Alphafold v.3 (Figure 4). Interestingly, we revealed four different states of the S_1_M_2_ complex, which feature different angles between M-S-M subunits (Figure 4A). The intermediate state O_2_ was predicted only in the case of one protein sArdN interacts with the S-subunit (Figure 4B). Interestingly, that the state C_1_ was predicted in two cases: 2 x sArdC and 2 x sArdN interacting with the S-subunit and the structural alignment of the protein complex in the C_1_ state shows highly similar structure and the angles between M-S-M subunits (Figure 4C). A detailed view of the interactions of Ards with the S-subunit reveals that sArdC forms dimers (Figure 4D) and sArdN interacts as two separate monomers (Figure 4E). This is remarkable because two different agents sArdC and sArdN with different interaction pathways nevertheless showed the same conformational changes of the protein complex S_1_M_2_. This fact significantly increases the possibility of the existence of the EcoKI-ArdA complexes in the predicted C_1_ state in nature. For reference, other interactions (with DNA and a single Ards, etc) were also predicted (Figure S1).

**Figure 4.**
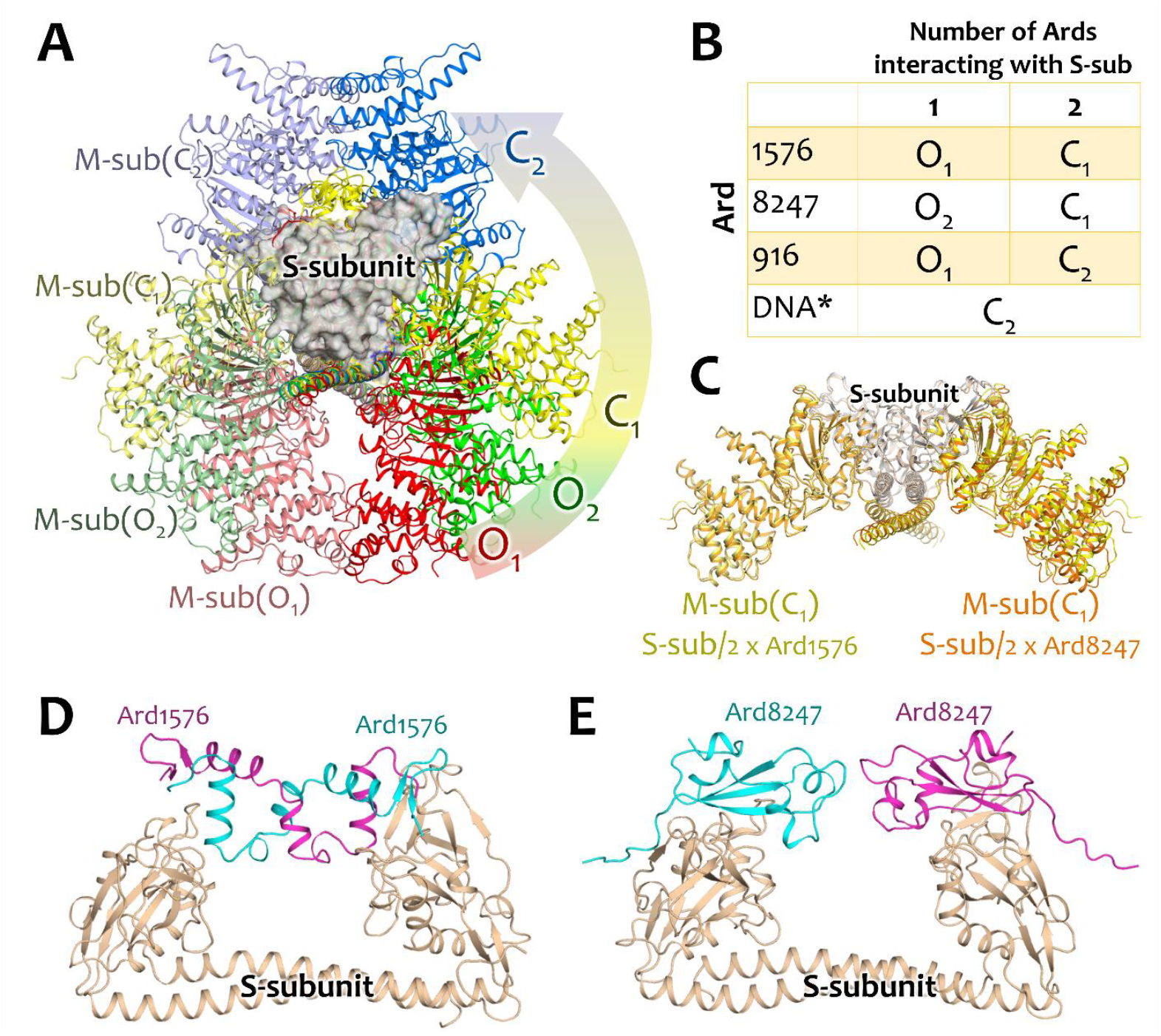
Structure predictions. **A -** scheme of EcoKI protein complex functioning. Two open (O_1_ and O_2_) and two closed (C_1_ and C_2_) states are distinguished. **B** – EcoKI states during interactions with different agents. ^*^DNA means dsDNA (AAAACACGTGTGTGCAA). **C** – Structural alignment of EcoKI with 2 x sArdC (ArdA_1576) and with 2 x sArdN (ArdA_8247). In both cases the intermediate state C_1_ is predicted. **D** – Interaction between the S-subunit of EcoKI with 2 x ArdA_1576 predicts that sArdC (ArdA_1576) forms dimers, and **E** – with 2 x sArdN (ArdA_8247) predicts that two separate sArdN (ArdA_8247) interact with the S-subunit as monomers.

Alpha fold predictions allowed us to assume that sArdN and sArdC proteins could work as an antirestriction agents. We confirmed this hypothesis in lambda phage experiments (figure 5). It was demonstrated that the sArdN and sArdC proteins exhibit «full-sized» antirestriction activity, similar to that of ArdA_R64.

**Figure 5.**
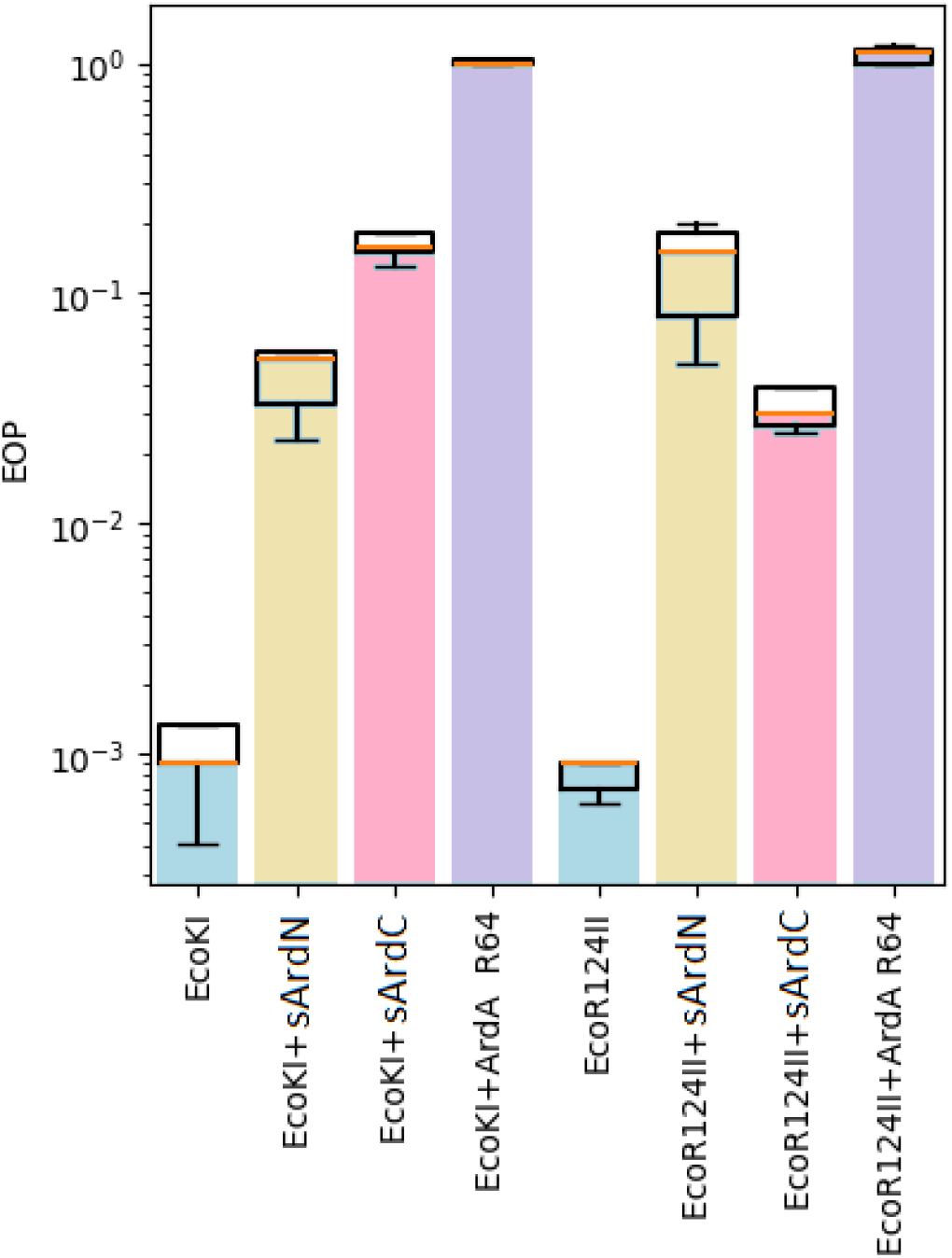
The results of the λ_0_ phage plaquing (EOP) on a lawn of *E. coli* cells containing genes of various RMI systems of gram-negative bacteria. EOP represents a ratio of a phage titer obtained on the experimental lawn relative to the TG1 lawn, which is sensitive to phage infect io n. Columns: EcoKI - TG1 pACYCEcoKI; EcoKI+sArdN – TG1 pACYCEcoKI, pIR-DPAl-ArdA-8247; EcoKI+sArdC – TG1 pACYCEcoKI, pIR-DPAl-ArdA-1576; EcoKI + ArdA_R64 – TG1 pACYCEcoKI, pIR-DPAl-ArdA-R64; EcoR124II - TG1 pKF650; EcoR124II+sArdN – TG1 pKF650, pIR-DPAl-ArdA-8247; EcoAI+sArdC – TG1 pKF650, pIR-DPAl-ArdA-1576; EcoR124II + ArdA_R64 – TG1 pKF650, pIR-DPAl-ArdA-R64.

As we can see from the data presented in Figure 5, the genes *sardN* and *sardC* have a certain specificity to different RMI systems. Thus, sArdC is more effective against EcoR124II, while sArdN is more effective against EcoKI. Additionally, we demonstrated that both studied *sard*-genes inhibit methylation of λ_0_ phage (Table 2).

**Table 2.** Data on the λ phage plating: unmethylated λ_0_ phage and methylated and λ_k_ phages (were infected with the phage λ_0_ and after the one-step growth cycle), the obtained phage were grown on *E. coli* AB1157 cells containing the complete EcoKI restriction-modification system. ArdA from conjugative plasmid R64 was used as a control.

| | AB1157 + $\lambda_k$ | AB1157 + $\lambda_{ksardC}$ | AB1157 + $\lambda_{ksardN}$ | AB1157 + $\lambda_{kR64}$ |
| --- | --- | --- | --- | --- |
| The number of phage colonies (-6 dilution)* | 15240±400 | 2760±70 | 2628±120 | 23±5 |
| Antimethylation rate | 1 | 1,8x10<sup>-1</sup> | 1,7x10<sup>-1</sup> | 1,5x10<sup>-3</sup> |

\*The results were averaged over three replicate experiments.

It is known that DNA mimetic proteins are able to specifically inhibit various DNA-binding proteins. For example, Ugi (from *Bacillus subtilis*) is an inhibitor of the uracil DNA glycosylase (Mol et al., 1995). Overall, the ability of these DNA-mimicking proteins to specifically inhibit different DNA-binding proteins makes them a promising tool for regulating a range of intracellular processes, including gene expression.

Here we described two totally new DNA-mimicking proteins which are twice smaller than classic ArdA proteins

## Discussion

The classical mechanism of new gene acquiring during evolution is considered to be duplication followed by the further mutagenesis of one of the copies (Magadum et al., 2013). In our work, we demonstrated that deletions of different parts of the *ardA* gene lead to the formation of two active gene variants encoding either the C-terminal or N-terminal regions of the full-length protein (Figures 1 and 2). Phylogenetic studies show that both the C-terminal and N-terminal small ArdA proteins (which we named sArdC and sArdN) are conserved throughout evolution and form homologous genes within certain bacterial taxa (Figure 3). As seen in Figures 3A and 3B, *sardC* genes cluster within taxa such as the *genus Lactococcus* and the phylum *Actinomycetes*. The fact that above the taxon level of Actinomycetes, *sArdC* and *sArdN* begin to cluster with their respective full-length ArdA genes strongly suggests repeated formation of small *ardA* genes in bacteria during evolution.

Notably, pairs of sArdA proteins derived from different sources exhibit specificity to different restriction-modification systems. For instance, sArdC is more effective against EcoR124II, while sArdN is effective against EcoKI. It has been suggested in previous work, that specificity of ArdA proteins to DNA-binding proteins may depend on an additional domain, as observed in *B. bifidum*(M. V Gladysheva-Azgari et al., 2023). However, our current findings support the idea that binding specificity of DNA-mimicking proteins to their targets could also be achieved by very short proteins. This effect opens up prospects for creating agents to specifically inhibit DNA-binding regulators.

Alphafold structure prediction allowed us to reveal the dynamics of the EcoKI-Ard protein complexes with four different states (two open – O_1_ and O_2_ and two closed – C_1_ and C_2_). Interestingly, that interaction with different agents (2 x sArdC and 2 x sArdN) provided the similar predictions of the EcoKI conformation (C_1_ state). This state, on the one hand, being intermediate between open O_1_ and closed C_2_ states (it shows the angle between M-S-M subunits approximately 120 degrees, Figure 4A), on the other hand the EcoKI complex is inhibited during this interaction as it is shown by the results of the λ_0_ phage plaquing (EOP) on a lawn of *E. coli* cells containing genes of various RMI systems of gram-negative bacteria (Figure 5), therefore, we named it a closed state C_1_. Thus, the possible presence of two distinct closed states of the EcoKI protein complex might indicate totally different molecular mechanisms of inhibiting of RM systems, although additional structural studies should be performed to directly prove our hypothesis.

Structural modeling revealed that sArdC from *L. cremoris* B-1576 is structurally homologous to the interface between subunits of the ArdA dimer. Previous research by Zavilgel’skii and Rastorguev demonstrated that mutagenesis of this interface sequence reduces anti-restriction activity of ArdA and completely eliminates its antimethylation capability (Rastorguev and Zavilgelsky, 2003). Evidently, sArdC 1576 retains the activity of this part of the antirestrictase structure, which appears to have high antimethylase activity. This hypothesis is supported by data presented in Table 2, showing that only about 15% of phage particles grown on r+m+ cells containing sArdC are methylated. Surprisingly, sArdN from *C. pilbarense* B-8247, which is homologous to the N-terminus of ArdA and theoretically cannot mimic the subunit interaction interface, is also highly efficient at antimethylation. This paradox awaits further investigation.

Finally, high antimethylation ability combined with relatively weak antirestriction could be beneficial for chromosomal genes during bacteriophage infection or plasmid conjugation and possible presence of different mechanisms of interaction pathways between Ards and RMI systems provides fundamental basis for future experiments.

## Funding

The work on searching and cloning antirestriction genes, antirestriction assay, phylogenetic studies were supported by RSF (project 24-74-00024).

The work on structural modeling and research was supported the Ministry of Science and Higher Education of the Russian Federation (agreement 075-03-2025-662, project FSMG-2025-0003).

The work on antimethylation measurements was carried out as part of a State Assignment for the Research Center Kurchatov Institute.

